# Whole genome assemblies of *Zophobas morio* and *Tenebrio molitor*

**DOI:** 10.1101/2022.12.21.521312

**Authors:** Sabhjeet Kaur, Sydnie A Stinson, George C diCenzo

## Abstract

*Zophobas morio* (=*Zophobas atratus*) and *Tenebrio molitor* are darkling beetles with industrial importance due to their use as feeder insects, their potential for use in aquafeed and human food products, and their apparent ability to biodegrade various plastic polymers. We report draft genome assemblies for *Z. morio* and *T. molitor* generated from Nanopore and Illumina data. Following scaffolding against published genomes, haploid assemblies of 462 Mb (scaffold N90 of 16.8 Mb) and 258 Mb (scaffold N90 of 5.9 Mb) were produced for *Z. morio* and *T. molitor*, respectively. Gene prediction led to the prediction of 28,544 and 19,830 genes for *Z. morio* and *T. molitor*, respectively. BUSCO analyses suggested both assemblies have a high level of completeness; 91.5% and 89.0% of the BUSCO endopterygota marker genes were complete in the *Z. morio* assembly and proteome, respectively, while 99.1% and 92.8% were complete in the *T. molitor* assembly and proteome, respectively. Phylogenomic analyses of four genera from the family Tenebrionidae yielded phylogenies consistent with those previously constructed based on mitochondrial genomes. Synteny analyses revealed large stretches of macrosynteny across the family Tenebrionidae, as well as numerous within-chromosome rearrangements. Finally, orthogroup analysis identified ∼28,000 gene families across the family Tenebrionidae, of which 8,185 were identified in all five of the analyzed species, and 10,837 were conserved between *Z. morio* and *T. molitor*. We expect that the availability of multiple whole genome sequences for *Z. morio* and *T. molitor* will facilitate population genetics studies to identify genetic variation associated with industrially relevant phenotypes.

## INTRODUCTION

The order Coleoptera is the largest order of the animal kingdom, accounting for 25% of known animal species and 40% of known insect species. Better known as beetles, the order Coleoptera consists of over 360,000 known species (Audisio *et al*. 2015), with some researchers estimating the true number of beetle species to be 1.5 million (Stork *et al*. 2015). Despite the taxonomic richness of this order, beetles are under-represented in genome sequencing projects. A recent meta-analysis noted that the phylum Arthopoda is the most under-sequenced animal phylum, with the order Coleoptera being the most under-sequenced order within the phylum (Hotaling *et al*. 2021a). A separate study found that the number of publicly available genome sequences for beetle species is five-fold less than expected given the total number of available insect genomes (Hotaling *et al*. 2021b). Significant efforts are therefore required to increase the number of available beetle genome sequences if the Earth BioGenome Project is to succeed (Lewin *et al*. 2018, 2022).

Tenebrionidae, commonly known as darkling beetles, is a family within the order Coleoptera that is estimated to contain ∼20,000 species (Bouchard *et al*. 2017). To date, the genomes of seven species within the family Tenebrionidae have been sequenced: *Tribolium castaneum* (Tribolium Genome Sequencing Consortium 2008; Herndon *et al*. 2020), *Tribolium madens, Tribolium freeman, Tribolium confusum, Asbolus verrucosus, Tenebrio molitor* (Eriksson *et al*. 2020; Eleftheriou *et al*. 2022), and *Zophobas morio* (= *Zophobas atratus*). The latter two species are of particular interest due to their industrial relevance. Commonly known as mealworms and superworms, respectively, the larvae of *T. molitor* and *Z. morio* are routinely used as feeder insects for pet reptiles and fish. They are also consumed by humans in some cultures (Ramos-Elorduy 2009), with additional societies becoming increasingly interested in their use in aquafeed and human food products as an alternative to other animal products (van Huis 2013; Ribeiro *et al*. 2018; Rumbos and Athanassiou 2021). Interestingly, several recent studies have provided evidence that mealworms and superworms have the potential to biodegrade various types of plastics (Yang *et al*. 2015a; Brandon *et al*. 2018; Yang *et al*. 2020; Peng *et al*. 2020). While the mealworm and superworm gut microbiota appear to play an important role in the breakdown of plastic polymers (Yang *et al*. 2015b; Peng *et al*. 2020), insect-encoded enzymes may also contribute to this process (Yang *et al*. 2021).

As the current study was in progress, high quality genome assemblies were made publicly available for *T. molitor* (Eleftheriou *et al*. 2022) and *Z. morio*, and annotations were provided for the *T. molitor* genome assembly. Here, we report independent genome assemblies and genome annotations for both *T. molitor* and *Z. morio*. The availability of genome annotations for both organisms will support future efforts to understand the mechanisms underlying plastic degradation by these insects. Likewise, the presence of multiple genome assemblies will facilitate population genetic studies that may be of value to breeders.

## MATERIALS AND METHODS

### Insect rearing and tissue collection

*T. molitor* and *Z. morio* larvae were purchased from a local pet store in Kingston, Ontario, Canada. The insect larvae were fed a diet of oatmeal, wheat bran, and carrot until harvest. For isolation of DNA, larvae were flash frozen in liquid nitrogen, and the head and legs of a single larvae per species were isolated for genomic DNA isolation. The collected tissues were crushed using sterile pestles in 1.5 ml tubes. For isolation of RNA, larvae were placed in 70% ethanol, following which their exoskeleton was cut open and the digestive tract removed. Digestive tracts were immediately flash frozen and stored at -80°C until use.

### DNA isolation and whole genome sequencing

For both *Z. morio* and *T. molitor*, crushed tissue from the head and legs of a single larvae was used for DNA extraction with Monarch Genomic DNA Purification Kits (New England Biolabs) following the manufacturer’s instructions. Briefly, insect tissue was resuspended in 10 μL Proteinase K, 20 μL EDTA, and 200 μL of the tissue lysis buffer supplied with the kit. The samples were incubated for three hours at 56°C with brief vortexing every 15 minutes, followed by treatment with RNase A for five minutes. Following binding of the genomic DNA to the silica membrane in the column, DNA was eluted with 100 µL of elution buffer and stored at -20°C until further processing.

Prior to Nanopore sequencing, the *Z. morio* DNA sample was size selected using a BluePippin instrument and a 0.75% agarose gel with the high pass protocol. The sample collected from the BluePippin instrument was cleaned-up and concentrated using an equal volume of AMPure XP Beads (Beckman Coulter), with the DNA eluted in in 60 μL DNAse and RNAse free water. Size selection was not performed for the *T. molitor* DNA sample. DNA samples were adjusted to a concentration of 20 ng/µL, following which library preparation was performed using a Ligation Sequencing kit (SQK-LSK100; Oxford Nanopore Technologies) following the manufacturer’s instructions. Sequencing was performed using a minION with R9.4.1 flow cells and the MinKNOW software. Basecalling was performed using GPU-enabled Guppy version 5.011+2b6dbffa5 and the high accuracy model (Oxford Nanopore Technologies).

Illumina sequencing and library preparation were performed at Génome Québec (Montréal, QC, Canada). Library preparation was performed using NxSeq AmpFREE Low DNA Fragment Library Kits (Lucigen) following the manufacturer’s instructions. Samples were then sequenced with 150 bp paired-end technology using a S4 flow cell on an Illumina NovaSeq 6000 instrument. Illumina reads were trimmed using platanus_trim version 1.0.7 (Kajitani *et al*. 2014).

### RNA isolation and sequencing

Total insect and microbial RNA was isolated from the digestive tracts of either one *Z. morio* or two *T. molitor* larvae using ZymoBIOMICS RNA miniprep kits (Zymo Research), following the manufacturer’s instructions.

RNA depletion, library preparation, and Illumina sequencing were performed at Génome Québec (Montréal, QC, Canada). Depletion of rRNA was performed using FastSelect kits (Qiagen) with probes from both the -rRNA fly and 5S/16S/23S rRNA kits, following the manufacturer’s instructions. However, the fly (*Drosphilia melanogaster*) rRNA probes were not efficient at removal of *Z. morio* or *T. molitor* rRNA as indicated by most of the sequencing reads mapping to beetle rRNA loci. Library preparation was completed using NEBNext Ultra II Directional RNA Library Prep kits (New England Biolabs), following the manufacturer’s instructions. Samples were then sequenced with 150 bp paired-end technology using a S4 flow cell on an Illumina NovaSeq 6000 instrument. Illumina reads were filtered using BBduk version 38.96 (Bushnell 2014) and then trimmed using trimmomatic version 0.39 (Bolger *et al*. 2014) with the following parameters: LEADING:3 TRAILING:3 SLIDINGWINDOW:4:15 MINLEN:36.

### Genome assembly

Nanopore reads were assembled into draft assemblies using Flye version 2.9-b1768 (Kolmogorov *et al*. 2019). Flye assemblies were polished once using Racon version 1.4.22 (Vaser *et al*. 2017) with Nanopore reads aligned to the draft assemblies with minimap2 version 2.20-r1061 (Li 2018 p. 2). Assemblies were further polished using Medaka version 1.4.1 (Oxford Nanopore Technologies) and the Nanopore reads. The assemblies were then polished using Pilon version 1.24 (Walker *et al*. 2014) with the trimmed Illumina reads mapped to the assemblies using bowtie2 version 2.4.4 (Langmead and Salzberg 2012 p. 2) and processed with samtools version 1.12 (Li *et al*. 2009). A final round of polishing was performed using Hapo-G version 1.2 (Aury and Istace 2021 p.) with the trimmed Illumina reads mapped to the assemblies using bowtie2.

In parallel, hybrid assemblies were constructed using MaSuRCA version 4.0.3 (Zimin *et al*. 2017) with the Nanopore and trimmed Illumina reads. The MaSuRCA assemblies were polished once using Pilon with the trimmed Illumina reads mapped to the assemblies with bowtie2. A second round of polishing was performed using Hapo-G with the trimmed Illumina reads mapped to the assemblies with bowtie2.

The Flye and MaSuRCA assemblies were merged using quickmerge version 0.3 (Chakraborty *et al*. 2016) and NUCmer version 4.0.0rc1 (Kurtz *et al*. 2004). Quickmerge was run using a minimum alignment length of 10,000 nucleotides and a length cutoff for anchor contigs approximately equal to the N50 of the self assembly. In addition, quickmerge was run twice for each insect species; once with the Flye assembly as the “self” assembly, and once with the Flye assembly as the “hybrid” assembly. The Flye and MaSuRCA assemblies were also merged using RagTag patch version 2.1.0 (Alonge *et al*. 2021) and NUCmer. As with quickmerge, RagTag patch was run twice for each insect species: once with the Flye assembly as the “query” sequence and once with the Flye assembly as the “target” sequence.

Haplotigs were purged from the Flye and MaSuRCA assemblies using purge_dups version 1.2.5 (Guan *et al*. 2020), with the self-self alignment performed using minimap2 version 2.18-r1015. The Flye and MaSuRCA assemblies purged of haplotigs were then merged using quickmerge and RagTag patch as described above. Purge_dups was then used to remove haplotigs from all merged assemblies created from either the full or purged assemblies. Finally, all *T. molitor* assemblies were scaffolded using RagTag scaffold and a previously published *T. molitor* assembly (European Nucleotide Archive [ENA] accession PRJEB44755) (Eleftheriou *et al*. 2022), whereas the *Z. morio* assemblies were scaffolded against a publicly available *Z. atratus* assembly (GenBank accession GCA_022388445.1).

Assembly statistics and measures of genome completeness and redundancy were performed for all assemblies as described below, and the results used to select one assembly to move forward. For *T. molitor*, the purged Flye assembly was selected for annotation. For *Z. morio*, the assembly chosen for annotation was the one created by using purge_dups on the assembly produced by running quickmerge with the purged MaSuRCA and the purged Flye assemblies as the hybrid and self assemblies, respectively.

The mitochondrial genomes of the *Z. morio* and *T. molitor* assemblies were identified by querying the assemblies using BLASTn version 2.10.1+ and previously published *Z. morio* (GenBank accession: MK140669.1) (Bai *et al*. 2019) or *T. molitor* (GenBank accession: KF418153.1) (Liu and Wang 2014) mitochondrial sequences, respectively. The searches revealed that scaffolding led to the *Z. morio* mitochondrion being merged with a nuclear DNA contig; the merged contig was therefore broken to isolate the mitochondrion as a single contig, and one copy of the overlap between the two ends of the mitochondrial contig were removed.

Submission of the genome assemblies to NCBI led to the identification of contamination in the assemblies, including the presence of adapter and microbial sequences. Contaminating sequences were removed from the *T. molitor* and *Z. morio* assemblies, and scaffolds were split at sites of contamination. Following decontamination, the nuclear genome was again scaffolded against the appropriate reference assembly as described above.

### Genome annotation

Low complexity regions of the genome were masked in a multi-step fashion. First, RepeatMasker version 4.1.2-p1 (Tarailo-Graovac and Chen 2009) was run on each assembly using the rmblast version 2.11.0 search engine (Tarailo-Graovac and Chen 2009), Tandem Repeats Finder version 4.0.9 (Benson 1999), species designation of “Insecta”, and the Dfam version 3.2 (Hubley *et al*. 2016) and RepBase RepeatMasker edition 20181016 (Bao *et al*. 2015) databases. Then, sdust version 0.1-r2 (Li 2018) was run on each assembly. Finally, for each assembly, regions masked by sdust but not masked by RepeatMaster were soft-masked in the fasta file returned by RepeatMasker using BEDtools version 2.26.0 (Quinlan and Hall 2010).

Following masking, the RNAseq reads were mapped to the genome assemblies using STAR version 2.7.10a (Dobin *et al*. 2013) and a two-pass procedure (Veeneman *et al*. 2015). A multi-step gene prediction was then performed using BRAKER version 2.1.6 (Hoff *et al*. 2016, 2019; Brůna *et al*. 2021). Gene prediction was first performed using BRAKER with the softmasking option and the RNAseq alignment file produced by STAR. A second, independent gene prediction was then performed using BRAKER with the softmasking option and a protein database consisting of i) the single-copy and multi-copy complete Endopterygota BUSCO genes identified in the assembly to be annotated; ii) proteins from the *T. castaneum* genome annotation (ENA accession PRJNA12540) (Tribolium Genome Sequencing Consortium 2008), and iii) proteins from a previously published *T. molitor* genome annotation (ENA accession PRJEB44755) (Eleftheriou *et al*. 2022). Subsequently the two gene prediction files were combined and filtered using TSEBRA version 1.0.3 (Gabriel *et al*. 2021) with default settings except for the intron_support parameter being set to 0.2. BRAKER dependencies included: samtools version 1.15-8-gbdc5bb8, bamtools version 2.5.2 (Barnett *et al*. 2011), the GeneMark suite version 4.69 (Lomsadze 2005; Lomsadze *et al*. 2014; Brůna *et al*. 2020), DIAMOND version 2.0.13 (Buchfink *et al*. 2015), AUGUSTUS version 3.4.0 (Stanke *et al*. 2006, 2008), and Spaln version 2.3.3d (Gotoh 2008; Iwata and Gotoh 2012 p. 2).

Following TSERBA, coding regions fully contained within another coding region on the same DNA strand were removed from the GFF annotation files. Likewise, the smaller of two overlapping genes on the same strand were removed. The GFF files were sorted and tidied using GenomeTools version 1.5.10 (Gremme *et al*. 2013), and converted to GenBank format with table2asn (NCBI). For both genome assemblies, protein fasta files containing all predicted isoforms were created by extracting the protein sequences from the GenBank files.

### Quality assessment of the genome assemblies

Genome assembly statistics were determined using the stats.sh function of BBmap version 38.90 (Bushnell 2014). Genome completeness metrics were calculated using BUSCO version 5.1.2 (Manni *et al*. 2021) with the eukaryota_odb10 and endopterygota_odb10 datasets with MetaEuk version 4-a0f584d (Levy Karin *et al*. 2020), HMMER version 3.2.1 (Eddy 2004), and BLAST+ version 2.12.0 (Camacho *et al*. 2009). Kmer QV was estimated using Yak version 4bdd51d (github.com/lh3/yak).

### Estimation of genome heterozygosity

The fastq files containing the forward and reverse Illumina sequencing reads were concatenated as a single file and used as input for Jellyfish version 2.3.0 (Marçais and Kingsford 2011) to count canonical kmer sizes with a size of 21 and create a kmer count histogram. The output of Jellyfish was passed to the GenomeScope webserver (qb.cshl.edu/genomescope/) (Vurture *et al*. 2017) to estimate genome heterozygosity.

### Comparative genomics

Orthofinder version 2.5.4 (Emms and Kelly 2015, 2019) was used to group Tenebrionidae proteins into orthogroups with the BLAST search engine and the -y option. As input, Orthofinder was provided proteins annotated in the two genome assemblies produced in this study (i.e., *T. molitor* and *Z. morio*), as well as the proteins annotated in the genome assemblies for *A. verrucosus* (GenBank accession GCA_004193795.1), *T. castaneum* (RefSeq accession GCF_000002335.3), *T. madens* (RefSeq accession GCF_015345945.1), and a previously published *T. molitor* genome (GenBank accession GCA_907166875.3). The other Tenebrionidae genomes available through National Center for Biotechnology (NCBI) Genome database were not included as they lacked annotations. Orthofinder dependencies included: BLAST version 2.10.1+, DendroBLAST (Kelly and Maini 2013), fastME version 2.1.4 (Lefort *et al*. 2015), MCL clustering (Van Dongen 2000), and the ETE tree library (Huerta-Cepas *et al*. 2016). A distance matrix of the proteomes was then computed using Jaccard distances and the “distance” function of the R package “philentropy” (Drost 2018), following which a dendogram was constructed using the “bionj” function of the R package “ape” (Popescu *et al*. 2012).

Synteny between Tenebrionidae genomes was detected and visualized using the D-Genies version 1.3.0 webserver (dgenies.toulouse.inra.fr) (Cabanettes and Klopp 2018). First, scaffolds larger than 1 Mb were extracted from each assembly of interest using pullseq version 1.0.2 (github.com/bcthomas/pullseq), and then sorted by length in descending order using seqkit version 2.2.0 (Shen *et al*. 2016). Pairwise analyses were then performed with D-Genies using the minimap2 version 2.24 aligner and the “many repeats” option. Dot plots were constructed for the *Z. morio* and *T. molitor* genome assemblies produced in the current work, previously published *Z. morio* (GenBank accession GCA_022388445.1) and *T. molitor* (GCA_907166875.3) genome assemblies, as well as genome assemblies for *T. madens* (GCA_015345945.1), *T. castaneum* (GCA_000002335.3), *T. freemani* (GCA_022388455.1), and *T. confusum* (GCA_019155225.1). The published genome assembly of *A. verrucosus* was excluded as all contigs were shorter than 1 Mb.

To detect SNPs between the *T. molitor* and *Z. morio* assemblies produced in this and previous studies, the assemblies were first aligned using NUCmer with the options “--minmatch 100 --mincluster 1000 --diagfactor 10 --banded --diagdiff 5”. The output of NUCmer was then fed into the show-snps function of the MUMmer package, run with the “-Clr” options.

### Phylogenomic analysis

A phylogenomic tree was constructed to show the evolutionary relationships between *T. molitor* (this study and GenBank accessions GCA_907166875.3 and GCA_014282415.2), *Z. morio* (this study and GCA_022388445.1), *T. madens* (GCA_015345945.1), *T. castaneum* (GCA_000002335.3), *T. freemani* (GCA_022388455.1), *T. confusum* (GCA_019155225.1), and *A. verrucosus* (GCA_004193795.1). First, highly conserved genes were detected in all genomes using BUSCO version 5.3.0 with the eukaryota_odb10 database. A set of 155 single copy orthologs present in all 10 genome assemblies were identified, following which each set of orthologs was individually aligned using MAFFT version 7.310 with the localpair option (Katoh and Standley 2013) and trimmed using trimAl version 1.4rev22 and the automated1 option (Capella-Gutiérrez *et al*. 2009). Trimmed alignments were then concatenated, and used as input to construct a maximum likelihood phylogeny with RAxML version 8.2.12 with the GAMMA model of rate heterogeneity and the JTT amino acid substitution model with empirical amino acid frequencies. The JTT model with empirical frequencies was chosen based on the results of a preliminary run of RAxML using automatic model selection. The final tree represents the bootstrap best true following 100 bootstrap replicates, and was visualized using the iTol webserver (Letunic and Bork 2016).

## RESULTS AND DISCUSSION

### *Z. morio* and *T. molitor* genome assemblies

Genomic DNA isolated from single *Z. morio* and *T. molitor* individuals was sequenced using a Nanopore minION (*Z. morio*: 2.79 Gb with a N50 read length of 14,531 nt; *T. molitor*: 8.84 Gb Gb with a N50 read length of 4,113 nt) and an Illumina NovaSeq to generate 150 bp paired-end reads (*Z. morio*: 220 Gb; *T. molitor*: 167 Gb). Multiple approaches to genome assembly were taken to increase the likelihood of producing a good-quality assembly (see Materials and Methods). Variations to the assembly pipeline included: i) performing either nanopore-only assembly using Flye or hybrid assembly using MaSuRCA, ii) optionally merging the Flye and MaSuRCA assemblies using quickmerge or RagTag, and iii) optionally purging haplotigs using purge_dups. Following scaffolding against existing genome assemblies (see Materials and Methods), a single assembly to move forward to annotation was chosen for each species based on the genome size, contig and scaffold N50, and BUSCO scores as a measure of assembly completeness and redundancy (**Table S1**).

Using the above approach, haploid genome representations of 462 Mb (scaffold N90: 16.8 Mb) and 258 Mb (scaffold N90: 5.9 Mb) were produced for *Z. morio* and *T. molitor*, respectively (**Table 1**). By comparison, the existing *Z. morio* and *T. molitor* genome assemblies are 478 Mb and 288 Mb, respectively, suggesting that some repetitive regions were collapsed in our assemblies. Alignment of the published *Z. morio* and *T. molitor* mitochondrial sequences (GenBank accessions MK140669.1 and KF418153.1) identified full-length mitochondrial genomes as single contigs in both assemblies.

**Table 1.** *Z. morio* and *T. molitor* genome assembly statistics.

|  | *Zophobas morio*<br>(This study) | *Zophobas morio*<br>(GCA_022388445.1) | *Tenebrio molitor*<br>(This study) | *Tenebrio molitor*<br>(GCA_907166875.3) |
| --- | --- | --- | --- | --- |
| Cumulative scaffold length (bp) | 461,985,688 | 478,145,070 | 258,289,333 | 287,839,991 |
| Number of scaffolds | 4,179 | 562 | 1,987 | 111 |
| Number of contigs | 8,884 | 937 | 6,142 | 167 |
| G+C content (%) | 33.34 | 33.44 | 36.17 | 36.72 |
| Number of Ns | 470,362 | 37,038 | 416,000 | 28,500 |
| Largest scaffold (Mb) | 74.58 | 78.287 | 32.201 | 33.043 |
| Scaffold N50 (Mb) | 48.007 | 53.051 | 20.827 | 21.886 |
| Scaffold L50 | 4 | 4 | 6 | 6 |
| Largest contig (Mb) | 1.58 | 37.237 | 2.345 | 20.298 |
| Contig N50 (Mb) | 0.228 | 6.738 | 0.205 | 6.327 |
| Contig L50 | 535 | 15 | 300 | 13 |
| BUSCO complete single-copy * | 1914 | 2008 | 2083 | 2101 |
| BUSCO complete multi-copy * | 30 | 26 | 22 | 10 |
| BUSCO fragmented * | 39 | 28 | 5 | 3 |
| BUSCO missing * | 141 | 62 | 14 | 10 |

\* Determined using the BUSCO endopterygota\_odb10 dataset.

Estimates of genome quality indicated that both the *Z. morio* and *T. molitor* assemblies are of high quality. Using Yak and the Illumina data, the accuracies of the haploid assemblies were estimated at QV36 and QV31 for the *Z. morio* and *T. molitor* assemblies, respectively. Additionally, 98.1% of the BUSCO endopterygota marker genes were identified as single-copy, complete genes in the *T. molitor* assembly (**Table 1**), which is comparable to the value of 98.4% in the existing *T. molitor* assembly. Likewise, 90.1% of the BUSCO endopterygota marker genes were identified as single-copy, complete genes in the *Z. morio* assembly (**Table 1**), which is slightly lower than the value of 94.5% in the existing *Z. morio* assembly.

### *Z. morio* and *T. molitor* genome annotations

Gene prediction was performed using the BRAKER pipeline (Hoff *et al*. 2016, 2019; Brůna *et al*. 2021), and a combination of gut RNA-seq data and a protein database (see Materials and Methods). This process resulted in 28,544 and 19,830 predicted genes for *Z. morio* and *T. molitor*, respectively (**Table 2**). In comparison, a previous study predicted 21,435 genes in the *T. molitor* genome (Eleftheriou *et al*. 2022); to date, there are no other reports of *Z. morio* gene predictions. BUSCO analyses of the proteomes indicated that the gene predictions are good quality; 89% and 93% of the endopterygota marker genes were identified in the *Z. morio* and *T. molitor* proteomes, respectively, compared to 96% in the previous *T. molitor* annotation (**Table 2**). Likewise, 94% and 95% of the eukaryota marker genes were identified in the *Z. morio* and *T. molitor* proteomes, respectively, compared to 96% in the previous *T. molitor* annotation (**Table S2**). The higher frequency of multi-copy BUSCO marker proteins in our proteomes likely reflects the inclusion of alternate isoforms, whereas alternate isoforms were not included in the previously-reported *T. molitor* annotation.

**Table 2.** *T. molitor* and *Z. morio* gene prediction statistics.

|  | *Zophobas morio*<br>(This study) | *Tenebrio molitor*<br>(This study) | *Tenebrio molitor*<br>(GCA_907166875.3) * |
| --- | --- | --- | --- |
| Number of genes | 28,544 | 19,830 | 21,435 |
| Number of transcripts | 30,408 | 21,405 | Not reported |
| Cumulative length of CDSs (bp) † | 34,142,077 | 26,175,824 | 25,230,147 |
| Median gene length (bp) ‡ | 1,087 | 1,319 | 1,147 |
| Median CDS length (bp) | 813 | 876 | 783 |
| Median exon length (bp) | 193 | 190 | NR |
| Median intron length (bp) | 54 | 54 | 55 |
| Number of intronless genes | 11,785 | 5,788 | 4,898 |
| Median number of exons per gene | 2 | 3 | 3 |
| Median number of exons per multi-exon gene | 5 | 5 | 4 |
| BUSCO complete single-copy ø | 1781 | 1897 | 2019 |
| BUSCO complete multi-copy ø | 108 | 75 | 15 |
| BUSCO fragmented ø | 61 | 47 | 21 |
| BUSCO missing ø | 174 | 105 | 69 |

\* Values in this column were taken from Eleftheriou *et al.* (2022), except for the BUSCO scores.
† The cumulative CDS length includes the length of all isoforms for the two genomes reported in this study.
‡ Does not include 5' or 3' untranslated regions (UTRs).
ø Determined using the BUSCO endopterygota\_odb10 dataset.

Gene summary statistics for *Z. morio* and *T. molitor* revealed several similarities. The median exon length, median intron length, median coding sequence (CDS) length, and median exon count per gene was similar in both species (**Table 2**). Likewise, the number of alternate transcripts per gene is comparable in the two species: ∼1.07 transcripts per gene in *Z. morio* compared to ∼1.08 transcripts per gene in *T. molitor*. On the other hand, the median gene length in our *T. molitor* assembly (1,319) is ∼21% longer than the median gene length (1,087) in our *Z. morio* assembly; however, this pattern does not hold true when comparing the median *Z. morio* gene length to the median gene length (1,147) in the previous *T. molitor* annotation. Additionally, whereas *Z. morio* encodes more genes than does *T. molitor*, the coding density of *T. molitor* (∼77 genes per Mb in our assembly) is higher than that of *Z. morio* (∼62 genes per Mb). Lastly, the percentage of genes lacking introns is higher in *Z. morio* (∼41%) relative to *T. morio* (between 23% and 29%, depending on the *T. morio* annotation) (**Table 2**).

Repetitive DNA and low complexity DNA was annotated in the *Z. morio* and *T. molitor* genome assemblies using RepeatMasker (Tarailo-Graovac and Chen 2009) and sdust (Li 2018), respectively (**Table 3**). Interspersed repeats accounted for 5.17% and 5.30% of the previously reported *Z. morio* and *T. molitor* genomes, respectively. The values were somewhat lower for our *Z. morio* and *T. molitor* assemblies at 4.38% and 3.88%, respectively, further indicating that some repetitive regions were collapsed in our assemblies. In general, the abundance of DNA transposons, long terminal repeat (LTR) elements, and long interspersed nuclear elements (LINEs) is similar between the two species although the precise make-up varies modestly (**Tables 3 and S3**). For example, when normalized by assembly length, *Tc1-IS630-Pogo* DNA transposons are ∼2-fold more abundant in *T. molitor*, whereas L2/CR1/Rex LINEs are ∼4-fold more abundant in *Z. morio*. Likewise, the abundance of helitrons is modestly different between species, with them being ∼2.7-fold more abundant in *Z. morio*. In contrast, the abundance of short interspersed nuclear elements (SINEs) differs dramatically and are >18-fold more abundant in *T. molitor* when normalized by assembly length (**Table 3**). Likewise, satellite sequences are >12-fold more abundant in *T. molitor* when normalized by assembly length (**Table 3**). The difference in satellite sequence abundance I s driven by the prevalence of the 142 bp TMSATE1 satellite sequence in *T. molitor* (Petitpierre *et al*. 1988; Davis and Wyatt 1989), which we detected 505 times in our *T. molitor* assembly and 415 times in the existing *T. molitor* assembly. On the other hand, low complexity DNA, as identified with sdust, is ∼6.4-fold more abundant in *Z. morio* compared to *T. molitor* (**Table 3**).

**Table 3.**
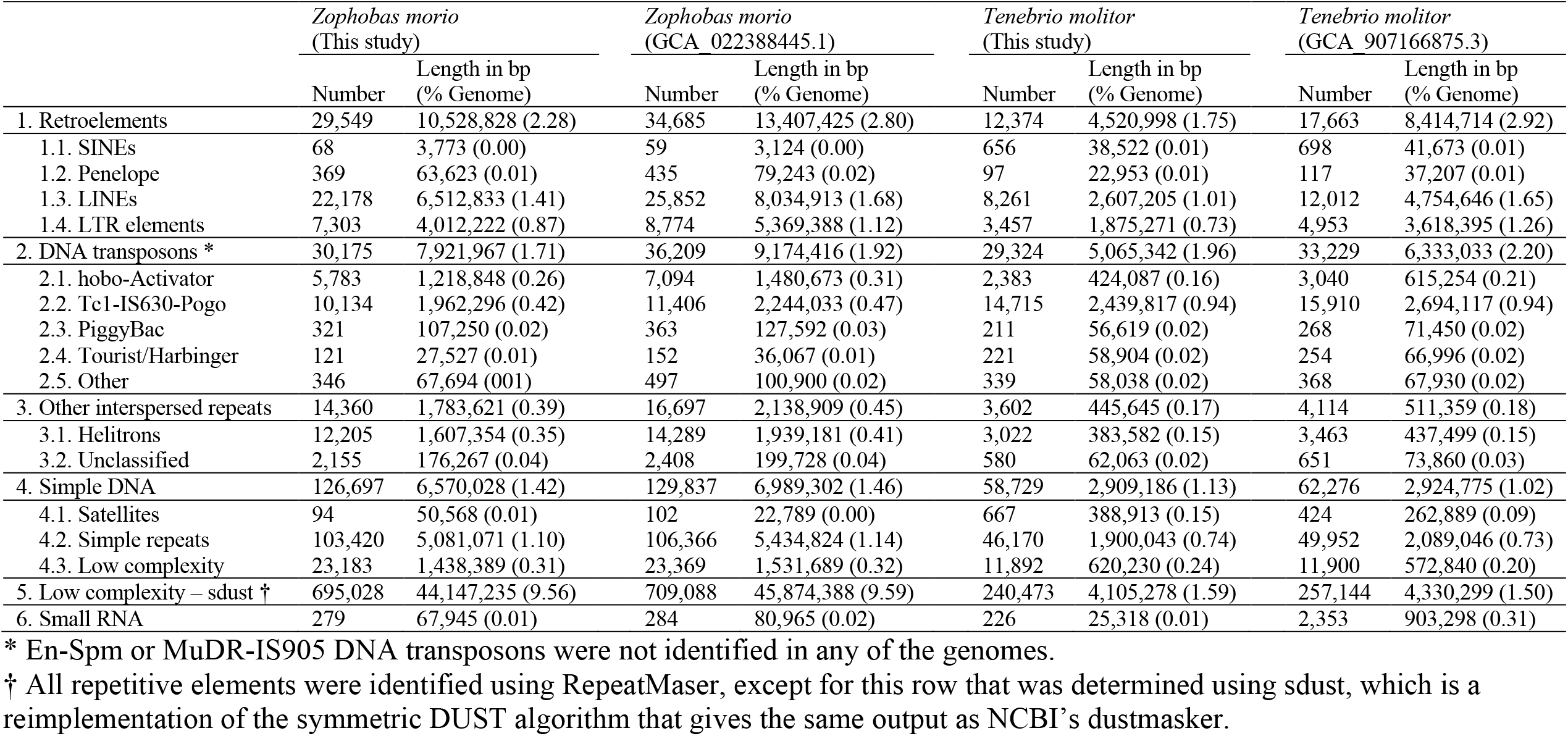
Repetitive elements identified in the *Z. morio* and *T. molitor* genome assemblies.

|  | *Zophobas morio*<br>(This study) |  | *Zophobas morio*<br>(GCA 022388445.1) |  | *Tenebrio molitor*<br>(This study) |  | *Tenebrio molitor*<br>(GCA 907166875.3) |  |
| --- | --- | --- | --- | --- | --- | --- | --- | --- |
|  | Number | Length in bp<br>(% Genome) | Number | Length in bp<br>(% Genome) | Number | Length in bp<br>(% Genome) | Number | Length in bp<br>(% Genome) |
| 1. Retroelements | 29,549 | 10,528,828 (2.28) | 34,685 | 13,407,425 (2.80) | 12,374 | 4,520,998 (1.75) | 17,663 | 8,414,714 (2.92) |
| 1.1. SINEs | 68 | 3,773 (0.00) | 59 | 3,124 (0.00) | 656 | 38,522 (0.01) | 698 | 41,673 (0.01) |
| 1.2. Penelope | 369 | 63,623 (0.01) | 435 | 79,243 (0.02) | 97 | 22,953 (0.01) | 117 | 37,207 (0.01) |
| 1.3. LINEs | 22,178 | 6,512,833 (1.41) | 25,852 | 8,034,913 (1.68) | 8,261 | 2,607,205 (1.01) | 12,012 | 4,754,646 (1.65) |
| 1.4. LTR elements | 7,303 | 4,012,222 (0.87) | 8,774 | 5,369,388 (1.12) | 3,457 | 1,875,271 (0.73) | 4,953 | 3,618,395 (1.26) |
| 2. DNA transposons * | 30,175 | 7,921,967 (1.71) | 36,209 | 9,174,416 (1.92) | 29,324 | 5,065,342 (1.96) | 33,229 | 6,333,033 (2.20) |
| 2.1. hobo-Activator | 5,783 | 1,218,848 (0.26) | 7,094 | 1,480,673 (0.31) | 2,383 | 424,087 (0.16) | 3,040 | 615,254 (0.21) |
| 2.2. Tc1-IS630-Pogo | 10,134 | 1,962,296 (0.42) | 11,406 | 2,244,033 (0.47) | 14,715 | 2,439,817 (0.94) | 15,910 | 2,694,117 (0.94) |
| 2.3. PiggyBac | 321 | 107,250 (0.02) | 363 | 127,592 (0.03) | 211 | 56,619 (0.02) | 268 | 71,450 (0.02) |
| 2.4. Tourist/Harbinger | 121 | 27,527 (0.01) | 152 | 36,067 (0.01) | 221 | 58,904 (0.02) | 254 | 66,996 (0.02) |
| 2.5. Other | 346 | 67,694 (0.01) | 497 | 100,900 (0.02) | 339 | 58,038 (0.02) | 368 | 67,930 (0.02) |
| 3. Other interspersed repeats | 14,360 | 1,783,621 (0.39) | 16,697 | 2,138,909 (0.45) | 3,602 | 445,645 (0.17) | 4,114 | 511,359 (0.18) |
| 3.1. Helitrons | 12,205 | 1,607,354 (0.35) | 14,289 | 1,939,181 (0.41) | 3,022 | 383,582 (0.15) | 3,463 | 437,499 (0.15) |
| 3.2. Unclassified | 2,155 | 176,267 (0.04) | 2,408 | 199,728 (0.04) | 580 | 62,063 (0.02) | 651 | 73,860 (0.03) |
| 4. Simple DNA | 126,697 | 6,570,028 (1.42) | 129,837 | 6,989,302 (1.46) | 58,729 | 2,909,186 (1.13) | 62,276 | 2,924,775 (1.02) |
| 4.1. Satellites | 94 | 50,568 (0.01) | 102 | 22,789 (0.00) | 667 | 388,913 (0.15) | 424 | 262,889 (0.09) |
| 4.2. Simple repeats | 103,420 | 5,081,071 (1.10) | 106,366 | 5,434,824 (1.14) | 46,170 | 1,900,043 (0.74) | 49,952 | 2,089,046 (0.73) |
| 4.3. Low complexity | 23,183 | 1,438,389 (0.31) | 23,369 | 1,531,689 (0.32) | 11,892 | 620,230 (0.24) | 11,900 | 572,840 (0.20) |
| 5. Low complexity – sdust † | 695,028 | 44,147,235 (9.56) | 709,088 | 45,874,388 (9.59) | 240,473 | 4,105,278 (1.59) | 257,144 | 4,330,299 (1.50) |
| 6. Small RNA | 279 | 67,945 (0.01) | 284 | 80,965 (0.02) | 226 | 25,318 (0.01) | 2,353 | 903,298 (0.31) |

\* En-Spm or MuDR-IS905 DNA transposons were not identified in any of the genomes.
† All repetitive elements were identified using RepeatMaser, except for this row that was determined using sdust, which is a reimplementation of the symmetric DUST algorithm that gives the same output as NCBI's dustmasker.

### *Z. morio* and *T. molitor* genome sequence variation

Using the k-mer-based approach of Genomescope together with Illumina data generated from a single individual per species, genome-wide heterozygosity was estimated to be 0.93% and 0.94% in *Z. morio* and *T. molitor*, respectively. By comparison, a higher heterozygosity of 1.43% was previously reported for *T. molitor* (Eleftheriou *et al*. 2022), which could be a result of the individuals coming from different populations. In comparing our *T. molitor* assembly with that of Eleftheriou *et al*. (2022), we identified ∼2.68 million SNPs, corresponding to ∼1.0% of the genomes. The between individual sequence variability was lower for the two *Z. morio* genomes; ∼2.39 million SNPs were identified, corresponding to ∼0.5% of the genomes.

### Comparative genomics of the family Tenebrionidae

A set of 155 single-copy orthologs was used to construct a maximum likelihood phylogeny to examine the evolutionary relationships between the seven Tenebrionidae species with sequenced genomes, representing four genera. The analysis suggests that the genera *Zophobas* and *Trilobium* are more closely related to each other than either are to the genus *Tenebrio*, and that the genus *Asbolus* is the most distantly related genus (**Figure 1**). These observations are consistent with a previously presented phylogeny constructed from 13 mitochondrial genes (Bai *et al*. 2019).

**Figure 1.**
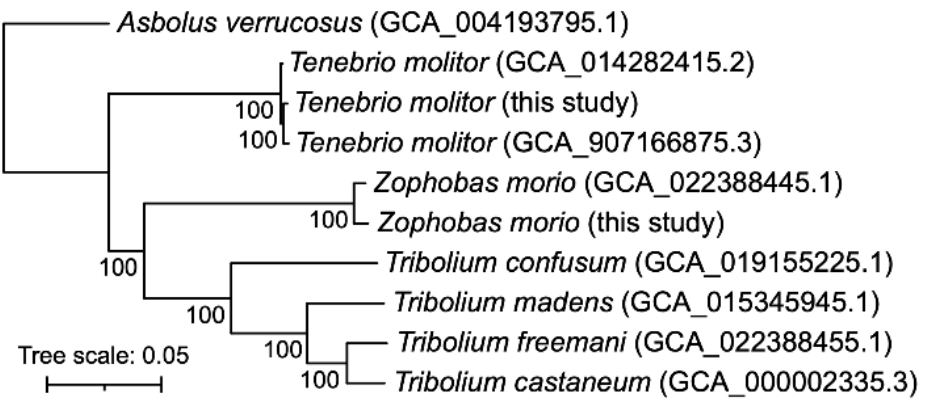
Phylogeny of the family Tenebrionidae. An unrooted maximum likelihood phylogeny of the family Tenebrionidae is shown. The phylogeny is drawn with *Asbolus verrucosus* as the outgroup based on the rooted species tree returned by OrthoFinder. The phylogeny represents the bootstrap best tree following 100 bootstrap replicates, which was prepared using RAxML with the concatenated alignment of 155 single-copy orthologs encoded by all 10 genomes. Values at the nodes represent the bootstrap support, while the scale bar represents the mean number of amino acid substitutions per site.

Dot plot analyses revealed large stretches of macrosynteny across the *Z. morio* and *T. molitor* genome assemblies (**Figure 2**). While many within-chromosome rearrangements were evident, no translocations were detected. Likewise, macrosynteny and within-chromosome rearrangement was observed when comparing the *Z. morio* or *T. molitor* assemblies to the genomes of four *Tribolium* spp. (**Figure S1-S4**). In addition, the previously reported whole-chromosome translocation within the *T. confusum* lineage was observed (Smith 1952; Samollow *et al*. 1983) (**Figure S1-S4**).

**Figure 2.**
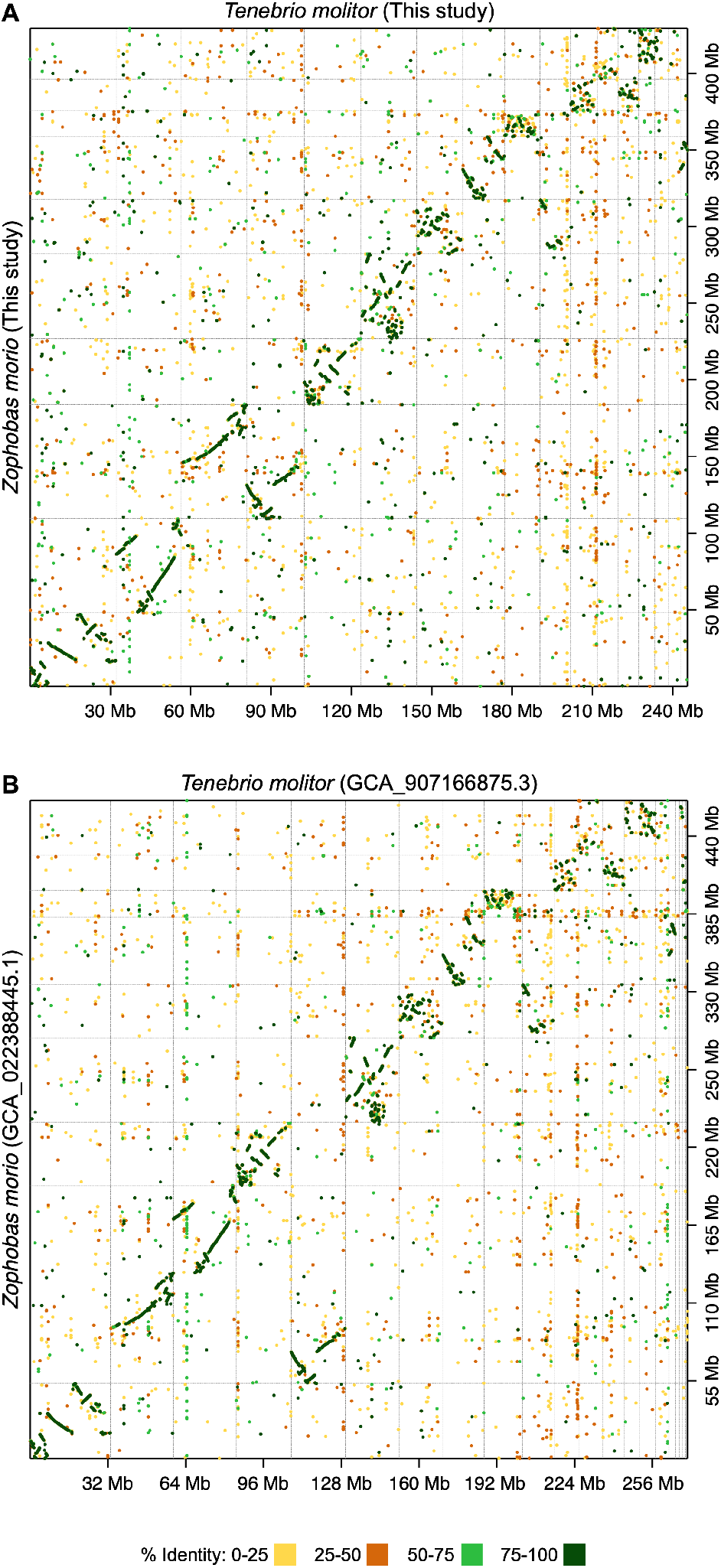
Macrosynteny between the *Zophobas morio* and *Tenebrio molitor* genomes. Dot plots are shown comparing the *Z. morio* and *T. molitor* genomes of (**A**) this study, or (**B**) previously published sequences (GCA_022388445.1 and GCA_907166875.3). Dot plots were created using D-Genies with the minimap2 aligner. Prior to the dot plot analyses, genomes were filtered to remove scaffolds less than 1 Mb in length for visualization purposes. Dashed grey lines delineate scaffolds. The dot colours indicate the average percent identity of the match.

To explore genome evolution within the family Tenebrionidae, OrthoFinder was used to group proteins from the five sequenced and annotated species of the genera *Asbolus, Tenebrio, Zophobas*, and *Tribolium*. This analysis identified a core set of 7,738 gene families present in all of the annotated genomes (**Figures 3 and S5**); this core set increases to 8,185 when including gene families present in at least one of the two *T. molitor* genome assemblies. The remaining 19,692 gene families were variably present in each genome, of which 14,761 gene-families (53% of all gene families) were species specific. In addition, a total of 10,837 gene families were conserved across *T. molitor* and *Z. morio*, when accounting for genes found in at least one *T. molitor* genome. Finally, a neighbour-joining tree constructed from a distance matrix of gene family presence/absence indicated that genome size was better correlated with proteome similarity than was phylogenetic relatedness (**Figure 3**).

**Figure 3.**
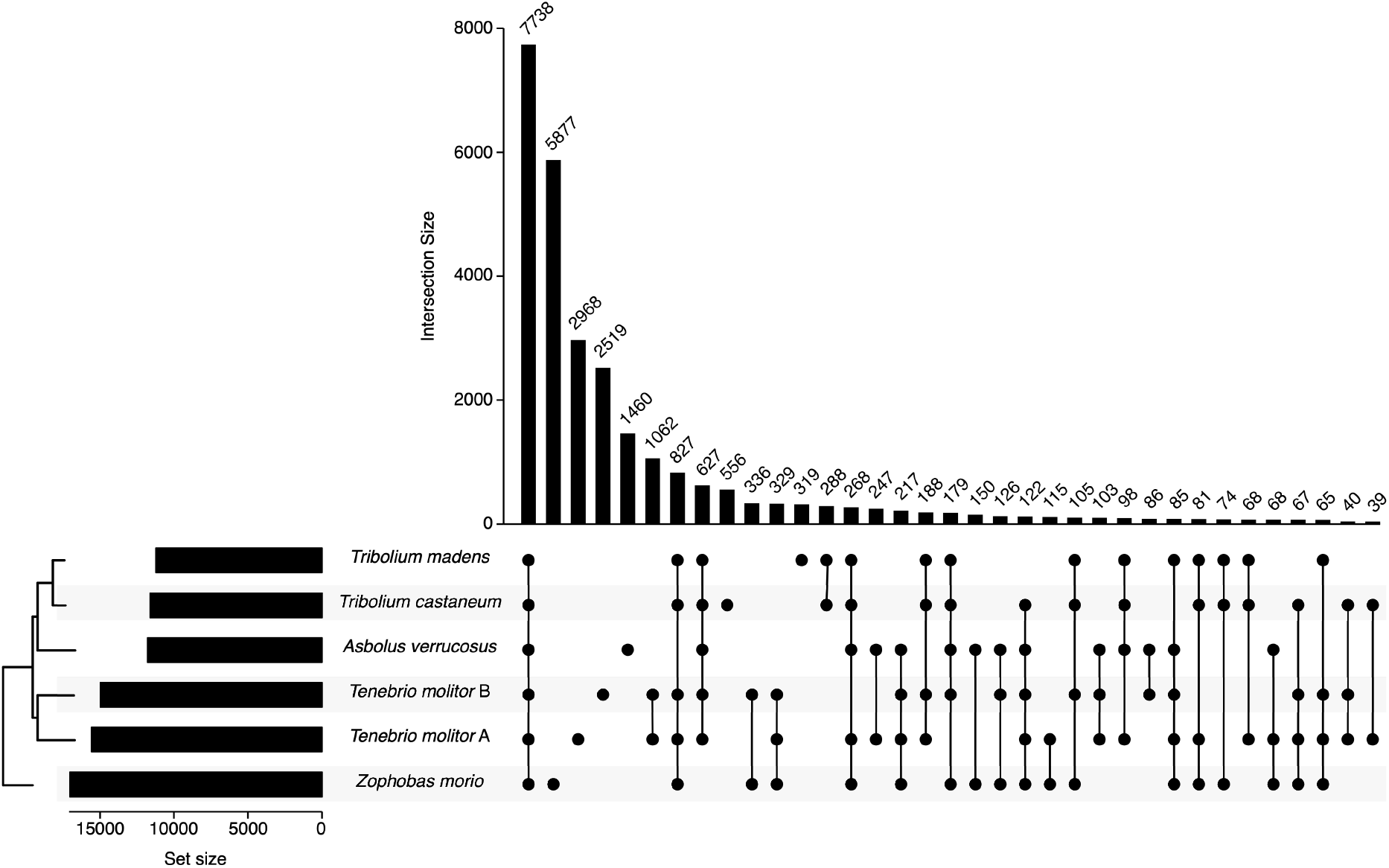
Conservation of gene families across the family Tenebrionidae. Orthofinder was used to group the annotated proteins of *A. verrucosus* (GenBank accession GCA_004193795.1), *T. castaneum* (RefSeq accession GCF_000002335.3), *T. madens* (RefSeq accession GCF_015345945.1), *T. molitor* A (GenBank accession GCA_907166875.3), *T. molitor* B (this study), and *Z. morio* (this study) into gene families. Gene family conservation was summarized using UpSetR (Conway *et al*. 2017), and the 35 most abundant intersections are shown (see **Figure S5** for a version with all intersections). The set size shows the total number of gene families in a given proteome, while the intersect size shows the number of gene families conserved across the indicated proteomes. The midpoint rooted dendogram presented in the bottom left represents the clustering of the proteomes creating using a neighbour-joining approach and a distance matrix constructed from the gene family presence/absence data and Jaccard distances.

## CONCLUSIONS

We report whole genome sequences for *Z. morio* (462 Mb; scaffold N90: 16.8 Mb) and *T. molitor* (258 Mb; scaffold N90: 5.9 Mb). The *Z. morio* genome is ∼1.8-fold larger than the *T. molitor* genome, in part due to an increase in interspersed repeats (20 - 25 Mb versus 10 - 15 Mb) and low complexity DNA (∼45 Mb versus ∼4 Mb), but also due to an increase in protein coding genes (28,544 versus 19,830 based on our gene predictions). Although many genomic rearrangements were detected, macrosynteny was observed between the *Z. morio* and *T. molitor* genomes, and 10,837 gene families were identified in both *Z. morio* and *T. molitor*.

We expect that the availability of multiple whole genome sequences for *Z. morio* and *T. molitor* will help facilitate future population genetics studies to identify genetic variation associated with industrially relevant phenotypes. In addition, these genome sequences will support studies of the microbiomes of darkling beetles by facilitating the removal of contaminating host DNA during metagenomic studies.

## DATA AVAILABILITY

All sequencing data generated in this study have been deposited to the NCBI under BioProject PRJNA820846. The Nanopore and Illumina DNA sequencing reads are available through the Sequence Read Archive (SRA) with the accession numbers SRR18507645, SRR18507646, SRR18507647, and SRR18507648. The Illumina RNA sequencing reads are available through the SRA with the accession numbers SRR18735291 and SRR18735292. Genome assemblies and annotations are available through the NCBI Genome database with the accession numbers: JALNTZ000000000 and JALNUA000000000. Scripts to repeat the computational work reported in this manuscript are available at: github.com/diCenzo_Lab/006_2022_Tenebrionidae_genomes.

## ACKNOWLEDGEMENTS

We thank Zhengxin Sun for kindly performing BluePippin size selection of the genomic DNA isolated from *Z. morio*. We also thank Philippe Daoust and other personnel at Génome Québec for the advice on sequencing strategies and for performing all the Illumina sequencing reported in this study. This research was enabled, in part, through computational resources provided by Compute Ontario (computeontario.ca) and Digital Research Alliance of Canada (alliancecan.ca).

