## Supplementary Figures S1-S5 for "Whole genome assemblies of *Zophobas morio* and *Tenebrio molitor*"

**Supplementary Materials for**
Whole genome assemblies of *Zophobas morio* and *Tenebrio molitor*

Sabhjeet Kaur, Sydnie A Stinson, George C diCenzo

George diCenzo


**This PDF file includes:**

Figures S1 to S5
Legends for Tables S1-S3

**Other supplementary materials for this manuscript include the following:**
Tables S1-S3

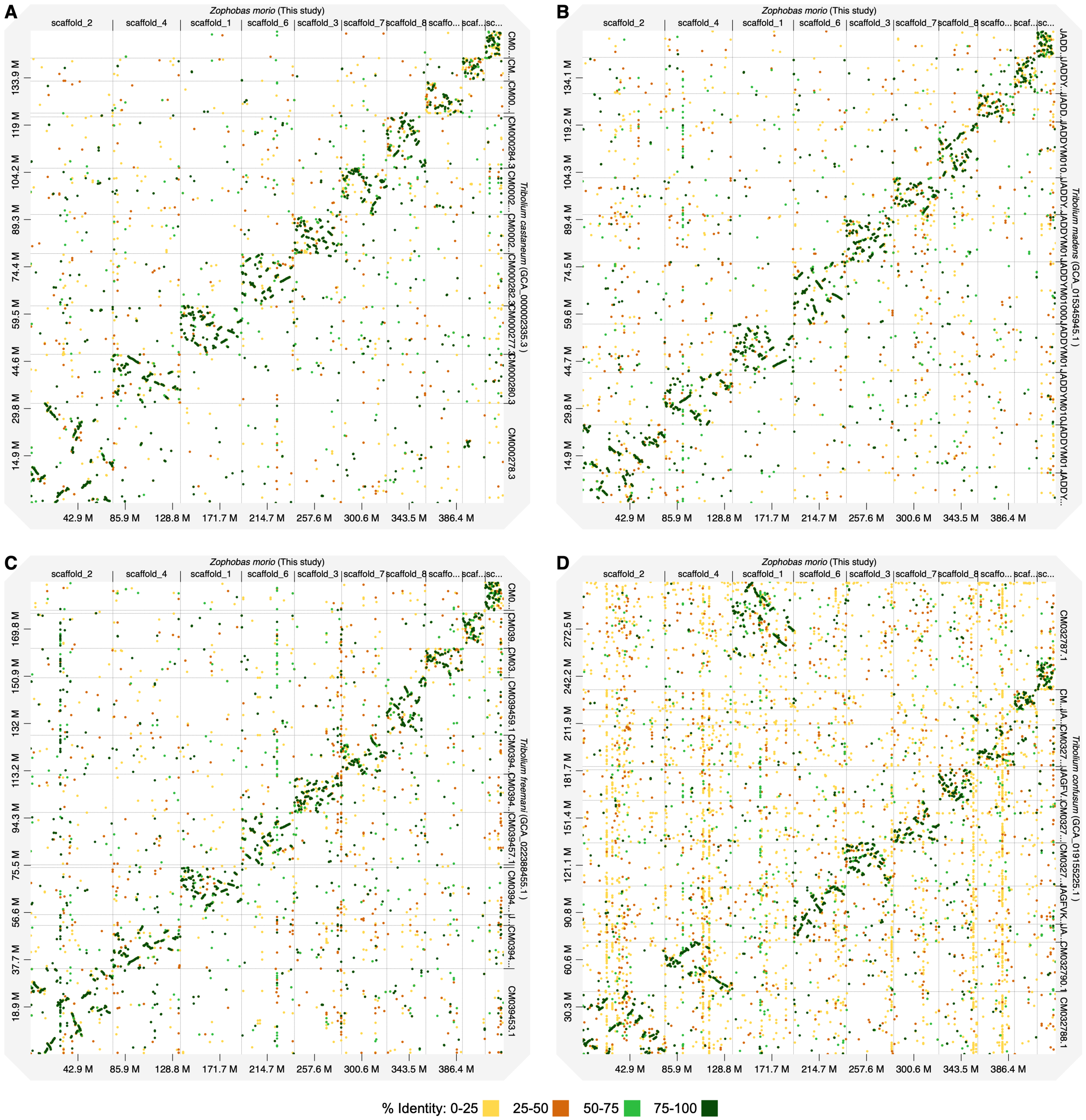


**Figure S1. Macrosynteny between *Zophobas morio* and *Tribolium* spp..** Dot plots are shown comparing the genomes of *Z. morio* (this study) and (**A**) *T. castaneum* (GCA_000002335.3), (**B**) *T. madens* (GCA_015345945.1), (**C**) *T. freemani* (GCA_022388455.1), and (**D**) *T. confusum* (GCA_019155225.1). Dot plots were created using D-Genies with the minimap2 aligner. Prior to the dot plot analyses, genomes were filtered to remove scaffolds less than 1 Mb in length for visualization purposes. Dashed grey lines delineate scaffolds. The dot colours indicate the average percent identity of the match.

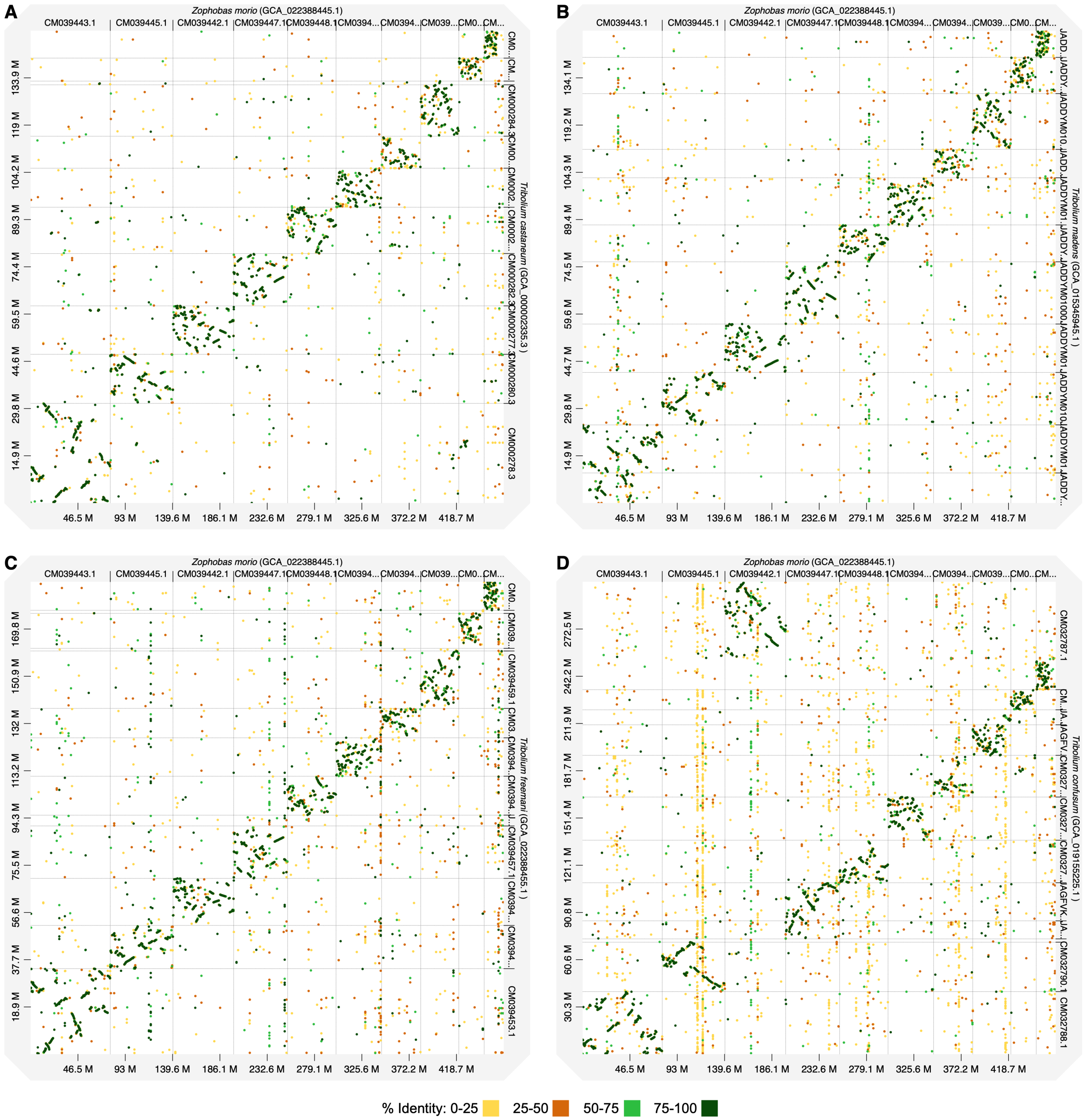


**Figure S2. Macrosynteny between *Zophobas morio* and *Tribolium* spp..** Dot plots are shown comparing the genomes of *Z. morio* (GCA_022388445.1) and (**A**) *T. castaneum* (GCA_000002335.3), (**B**) *T. madens* (GCA_015345945.1), (**C**) *T. freemani* (GCA_022388455.1), and (**D**) *T. confusum* (GCA_019155225.1). Dot plots were created using D-Genies with the minimap2 aligner. Prior to the dot plot analyses, genomes were filtered to remove scaffolds less than 1 Mb in length for visualization purposes. Dashed grey lines delineate scaffolds. The dot colours indicate the average percent identity of the match.

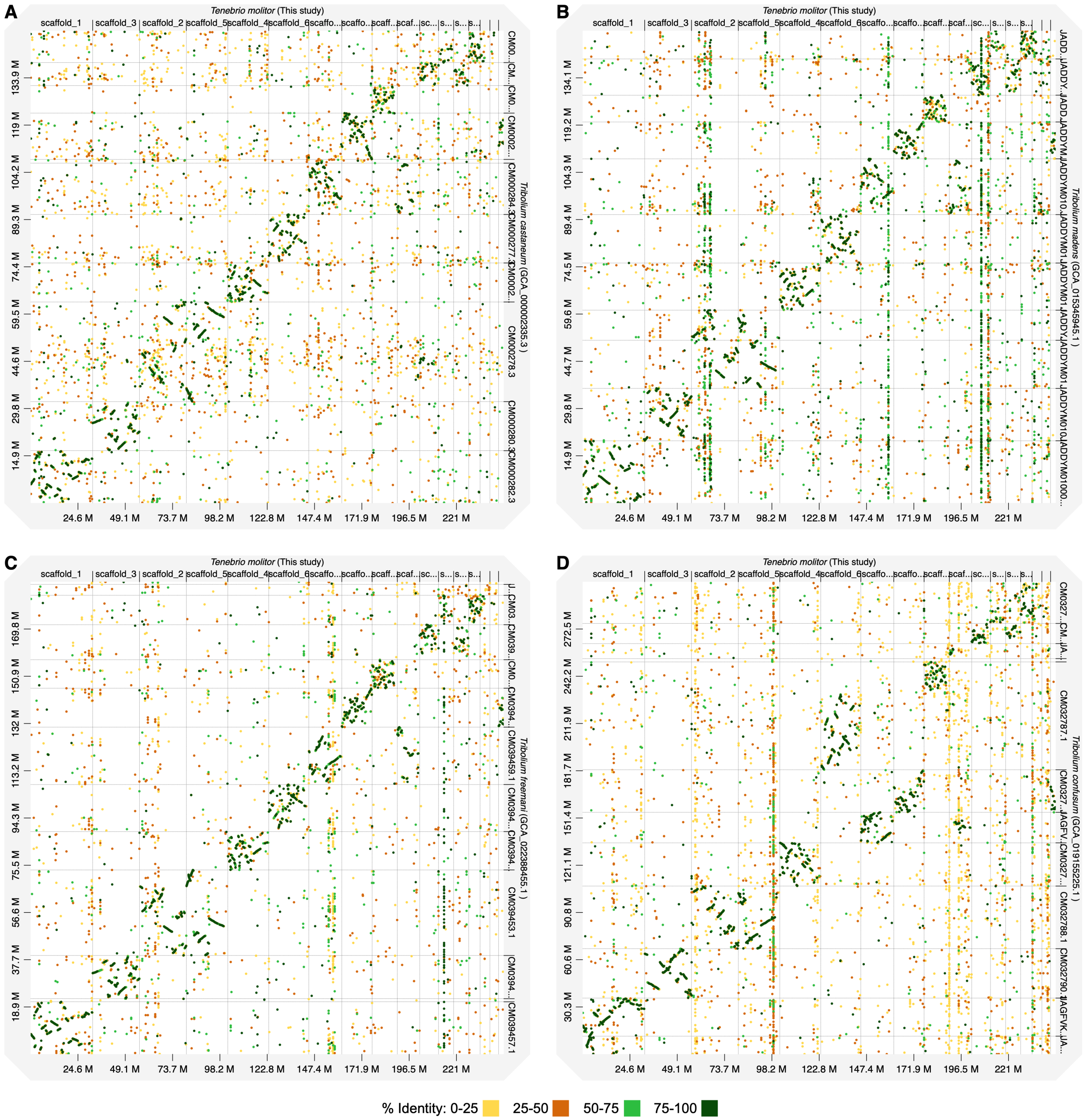


**Figure S3. Macrosynteny between *Tenebrio molitor* and *Tribolium* spp..** Dot plots are shown comparing the genomes of *T. molitor* (this study) and (**A**) *T. castaneum* (GCA_000002335.3), (**B**) *T. madens* (GCA_015345945.1), (**C**) *T. freemani* (GCA_022388455.1), and (**D**) *T. confusum* (GCA_019155225.1). Dot plots were created using D-Genies with the minimap2 aligner. Prior to the dot plot analyses, genomes were filtered to remove scaffolds less than 1 Mb in length for visualization purposes. Dashed grey lines delineate scaffolds. The dot colours indicate the average percent identity of the match.

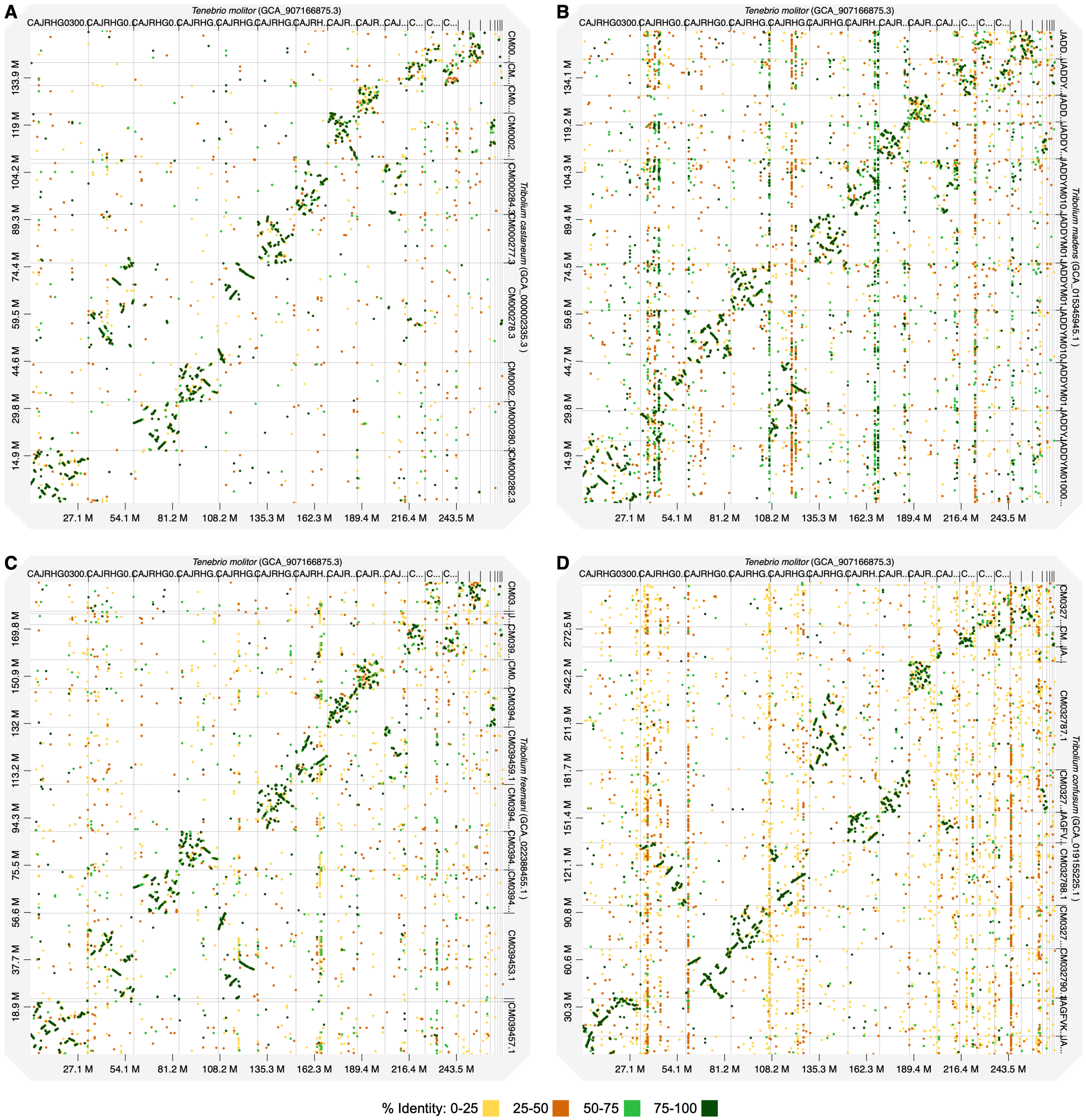


**Figure S4. Macrosynteny between *Tenebrio molitor* and *Tribolium* spp..** Dot plots are shown comparing the genomes of *T. molitor* (GCA_907166875.3) and (**A**) *T. castaneum* (GCA_000002335.3), (**B**) *T. madens* (GCA_015345945.1), (**C**) *T. freemani* (GCA_022388455.1), and (**D**) *T. confusum* (GCA_019155225.1). Dot plots were created using D-Genies with the minimap2 aligner. Prior to the dot plot analyses, genomes were filtered to remove scaffolds less than 1 Mb in length for visualization purposes. Dashed grey lines delineate scaffolds. The dot colours indicate the average percent identity of the match.

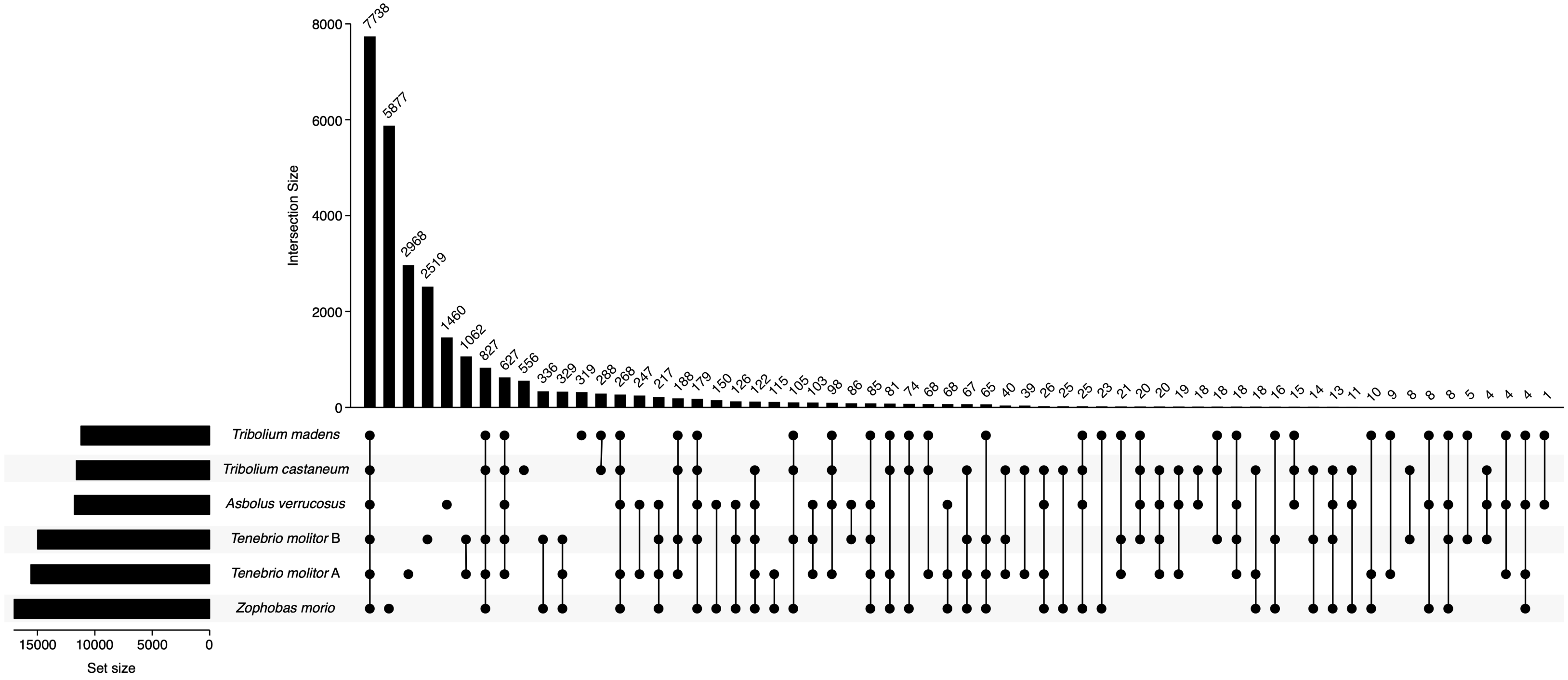


**Figure S5. Conservation of gene families across the family Tenebrionidae.** Orthofinder was used to group the annotated proteins of *A. verrucosus* (GenBank accession GCA_004193795.1), *T. castaneum* (RefSeq accession GCF_000002335.3), *T. madens* (RefSeq accession GCF_015345945.1), *T. molitor* A (GenBank accession GCA_907166875.3), *T. molitor* B (this study), and *Z. morio* (this study) into gene families. Gene family conservation was summarized using UpSetR. The set size shows the total number of gene families in a given proteome, while the intersect size shows the number of gene families conserved across the indicated proteomes.

### LEGENDS FOR SUPPLEMENTARY TABLES

**Table S1.** Assembly statistics, including BUSCO scores, for 20 *Zophobas morio* and 20 *Tenebrio molitor* genome assemblies are provided. The assembly selected to move forward to annotation is indicated in boldface font.

**Table S2.** Eukaryota BUSCO scores are reported for the new and previously published *Zophobas morio* and *Tenebrio molitor* genome assemblies and proteomes. Scores were calculated using the BUSCO eukaryota_odb10 dataset. No values are reported for the proteome of the previously published *Z. morio* genome as gene predictions were not performed for this genome.

**Table S3.** Classification of the LINEs and DNA transposons identified in the *Zophobas morio* and *Tenebrio molitor* genome assemblies.
